# Causal evidence of coherent theta rhythms in the modulation of multiregional brain communication

**DOI:** 10.1101/2023.09.20.558632

**Authors:** Gino Del Ferraro, Shaoyu Qiao, J. Isaac Sedillo, Bijan Pesaran

## Abstract

Neuromodulatory interventions seek to treat neuropsychiatric disorders by manipulating multiregional communication across the mesolimbic mood network. Modulations of multiregional communication are rarely measured directly and are often inferred from correlated neural activity such as neural coherence. Whether and how neural coherence reflects dynamic multiregional communication remains unclear. To address this limitation, we performed a causal-correlation analysis of theta-frequency (4-10 Hz) rhythms and mesolimbic multiregional communication. Selectively stimulating sender sites while recording from receiver sites revealed a mechanism of dynamic multiregional communication involving theta-coherent neural dynamics across a network of sender-receiver-modulator sites. Modulator site activity was highly theta-coherent with the receiver site activity, less theta-coherent with sender site activity, site specific and not shared by neighboring sites in the same region. These results reveal fundamental mechanisms of dynamic multiregional communication and support the use of theta-coherence as a target for neuromodulatory interventions in the mesolimbic mood network.

## Introduction

Multiregional communication occurs when the activity of neurons in one brain region, the sender, influences the activity of neurons in other regions of the brain, the receivers. Theta rhythms, periodic neural processes that repeat every 100-250 ms (4–10 Hz), are widely associated with multiregional communication. In sensory-perceptual systems, theta cycles chunk sensory processing to package information and transfer it from one region to another in order to support sensory processing and attention^1–4^. In hippocampal, occipital, prefrontal, striatal and orbitofrontal regions communication supports visual and spatial memory, decision making and reward learning^5–9^. The successful control of flexible behavior depends on regulating the flow of information between sender and receiver populations in cooperating regions of the cerebral cortex, basal ganglia and thalamus^10^ and so relies on the dynamic control of multiregional communication. Failures of communication between brain systems have also been implicated in a range of neuropsychiatric disorders that are amenable to treatment through neuromodulatory interventions, such as by deep brain stimulation (DBS)^11–13^.

Multiregional sender-receiver communication is most-often investigated through the analysis of correlated patterns of neural activity across different brain regions^2,14^. Correlated neural activity has been, for example, implicated in the coordination and temporal segregation of neural assemblies and dynamic routing of information flow due to modulatory context^15^. However, interpreting correlated patterns of neural activity in terms of sender-receiver communication channels can be unreliable, especially when putative sender-receiver neurons receive common inputs from other groups of neurons^16^. Common inputs can generate correlated patterns of neural activity in sender and receiver regions even when there is no influence of sender neuron activity on receiver neurons. Since common input is ubiquitous across pairs of nodes in large-scale brain networks^17^, and since theta rhythms are observed in neural activity across many regions of the brain^2^, the significance of correlated theta rhythms for multiregional communication remains uncertain.

Here, we use causal manipulations to more directly test the role that theta rhythms play in multiregional communication across the primate mesolimbic mood network. We electrically microstimulate and simultaneously record neural activity across the OFC, ACC, PCC, dPFC, motor cortex, somatosensory cortex, and posterior parietal cortex [PPC]) as well as the caudate nucleus [CN], putamen [Put], external globus pallidus [GPe], the amygdala (Amyg), the presubiculum [PrS], the substantia innominata [SI], the basal forebrain (BFB), and the nucleus accumbens (NAc). Simultaneously stimulating and recording across different brain regions allows us to identify pairs of sender-receiver sites without the confounding influence of common inputs. We then investigate the dynamic properties of correlated theta rhythms between the causally-identified sender-receiver sites and at other sites that do not respond to sender stimulation. By grounding the analysis of neural correlations in the results of causal manipulations, we provide new evidence for the role of theta rhythms in the dynamic control of multiregional communication.

## Results

### Identifying a multiregional communication channel

We delivered isolated, < 80 μA, electrical microstimulation currents while recording neural activity with an array of 200 microelectrodes in order to identify a multiregional network of sender-receiver (SR) communication channels (see **Methods**). We identified each communication channel by delivering a single, bipolar, biphasic microstimulation pulse at a given site, the sender, while simultaneously recording extracellular field potential responses at sites in the same and other brain regions, the receivers. The sampled multiregional network spanned 15 brain regions including seven cortical regions (orbital frontal cortex [OFC], anterior cingulate cortex [ACC], posterior cingulate cortex [PCC], dorsal prefrontal cortex [dPFC], motor cortex [M1], somatosensory cortex [S1], and posterior parietal cortex [PPC]) and eight subcortical regions (caudate nucleus [CN], putamen [Put], external globus pallidus [GPe], the amygdala (Amyg), the presubiculum [PrS], the substantia innominata [SI], the basal forebrain (BFB), and the nucleus accumbens (NAc). Responses to electrical stimulation were likely orthodromic due to direct connections to the stimulation site, eg OFC-CN, likely orthodromic due to indirect connections to the stimulation site, eg OFC-M1, or either antidromic responses on direct connections or due to indirect connections eg CN-OFC (**Table S1**). In each case, we interpreted the presence of a pulse of neural activity following each stimulation pulse as due to inter-regional communication across a communication channel.

**Figure 1a** presents an example communication channel. Stimulating a site in OFC, the sender site of the communication channel, reliably drove a response in M1, the receiver site, measured as a bipolar re-referenced stimulation-evoked LFP. In total, we tested 6910 pairs of electrodes across 132 pairs of brain regions. Of the 6910 tested pairs, we identified 110 sender-receiver channels spanning 21 region-pairs. For each of the 110 identified channels, examining the stimulation responses pulse-by-pulse revealed significant variability in how the receiver responded to sender stimulation. Figure 1b presents the average responses for the example OFC-M1 sender-receiver channel that took the form of “hit” events where a response was visible and “miss” events otherwise. Furthermore, “hit” and “miss” events were visible pulse-by-pulse and could be detected by an automated procedure trained and tested in a cross-validated manner (Fig 1c. see **Methods**).

**Figure 1.**
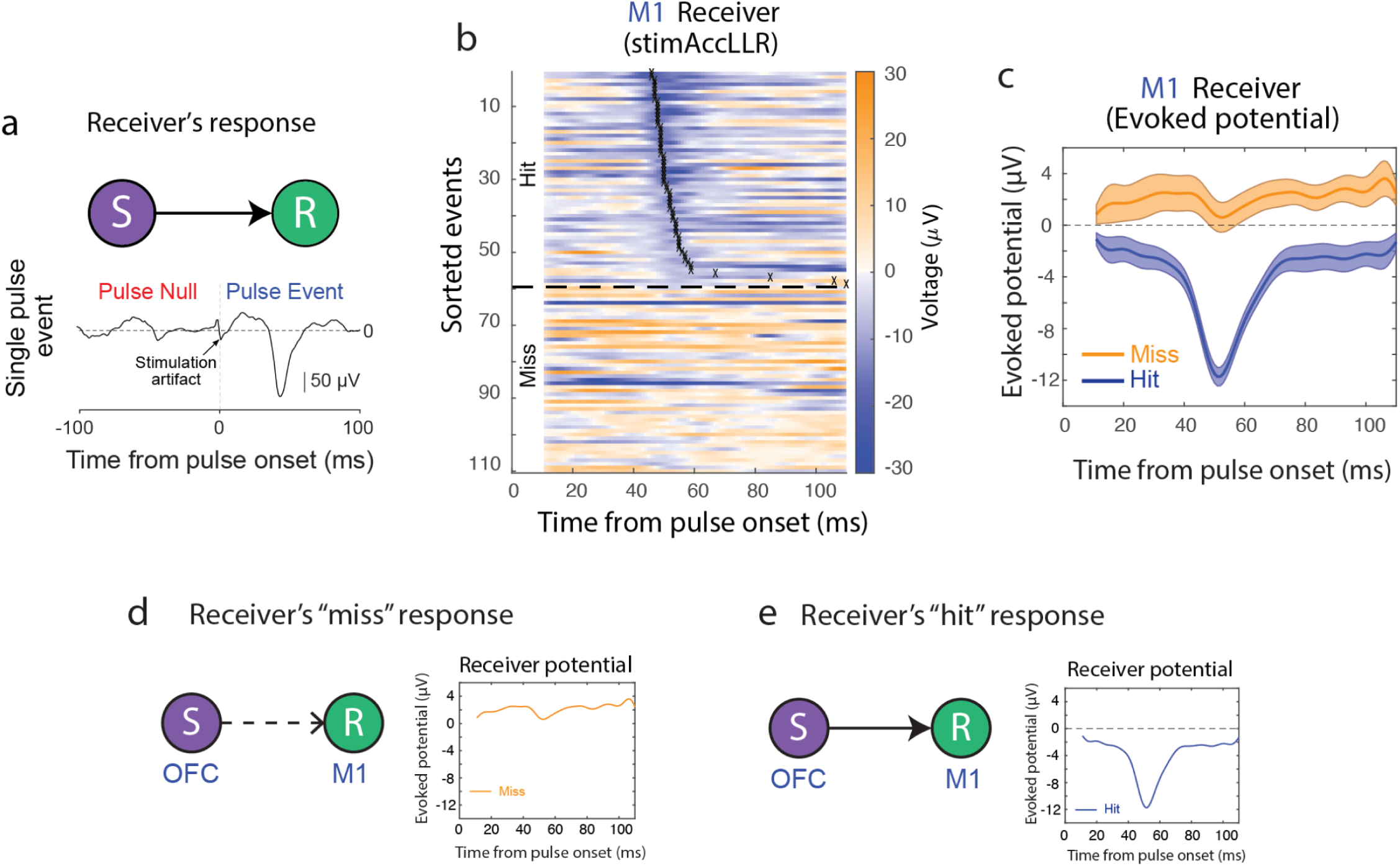
Sender-Receiver causal interaction. **(a)** Top: Sender-Receiver causal interaction diagram. Single, bipolar, biphasic, charge-balanced microstimulation pulses at the sender make it possible to causally identify sender-receiver communication. Bottom: An example of a receiver site response in CN brain region to a single microstimulation pulse at the OFC sender site (30 μs/phase). **(b)** Evoked M1 receiver responses sorted by pulse-response latency (crosses). **(c)** Pulse-triggered average evoked response for hit (blue) and miss events (orange). Shaded, ± SEM. **(d)** Microstimulation of the sender OFC site does not result in an evoked receiver response at the M1 site (miss event). **(e)** Microstimulation of the sender OFC site results in an evoked receiver response at the M1 site (hit event).

Broadly speaking, we interpret the absence of the expected stimulation responses as due to dynamic changes in neural excitability across a multiregional communication channel. When the excitability of neurons that support multiregional communication is low, electrical microstimulation is less likely to drive neurons to fire leading to absence of the expected stimulation response (Fig 1d). Conversely, when channel excitability is high, stimulation is more likely to lead to a response (Fig 1e).

### Multiregional communication and theta rhythms

We next asked how changes in SR communication may be associated with theta-frequency activity. Theta rhythms were clearly present across the sampled network, often occurring for a second or longer, and intermittent emphasizing their transient, dynamic nature (Fig 2a, gray shaded boxes). We asked whether the receiver response to sender stimulation, which was also dynamic, could be predicted from theta rhythms in the sampled theta network. Theta power immediately before stimulation at other sites in the network, termed modulator sites, significantly predicted changes in receiver responses immediately after stimulation. Figure 2b illustrates an example modulator site in M1 for an OFC-M1 communication channel. Examining single trials revealed variations in theta power on hit and miss trials (Fig 2b **middle**). For this example recording, when theta power at the modulator site was high, the receiver tended to respond to sender stimulation and channel excitability was high (Fig 2b **right**). Conversely, when theta power was low, the receiver tended not to respond to sender stimulation and channel excitability was low (Fig 2b **left**). To quantify this relationship, we performed an ideal observer analysis and tested whether theta power significantly predicted the presence or absence of the expected stimulation pulse (Fig 2c, **Methods**). For this example recording, increased theta power predicted an increased receiver response to sender stimulation with 72% accuracy (p-value corrected by false discovery rate for multiple comparison P_FDR_ = 3×10^-4^).

**Figure 2.**
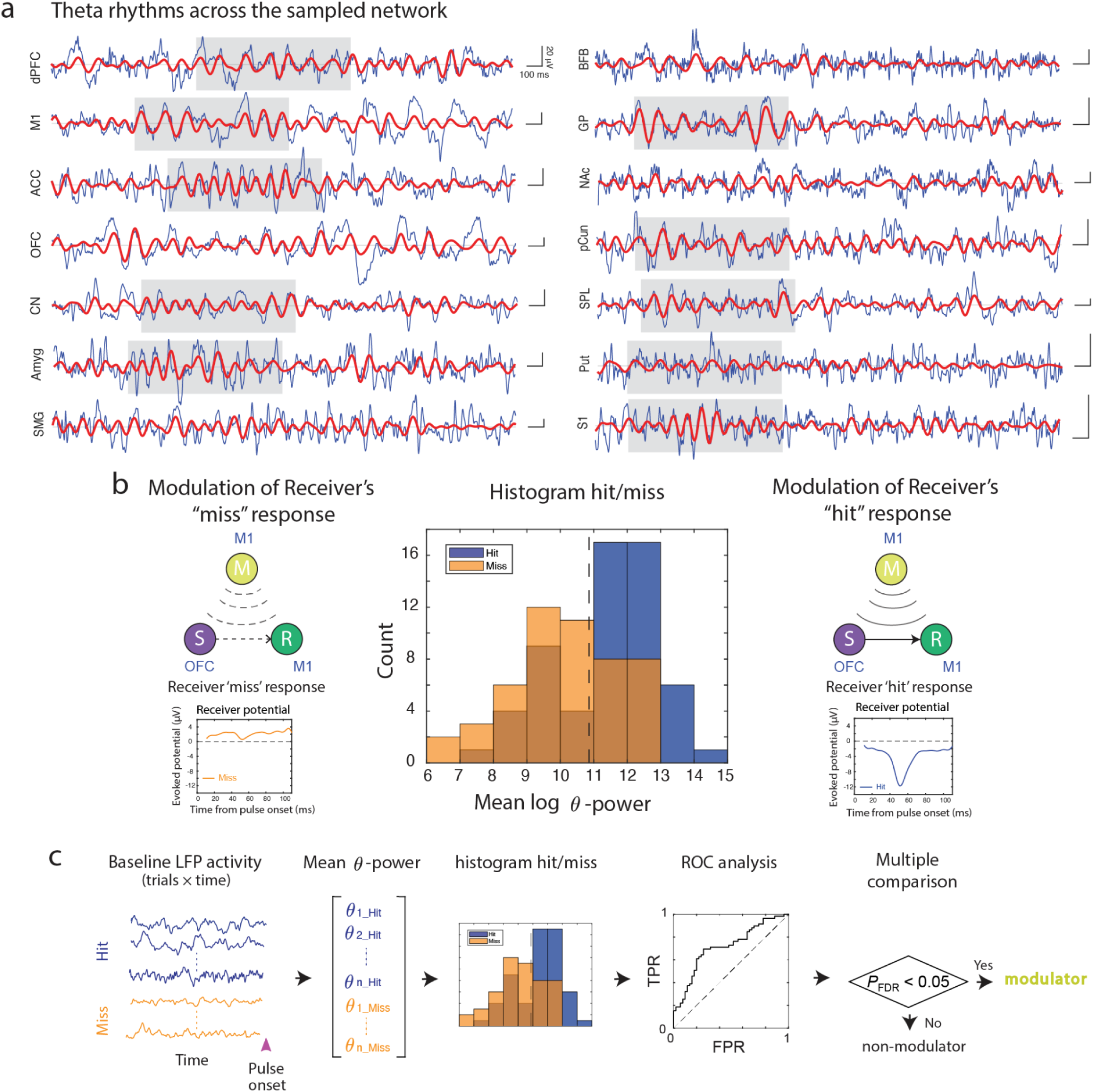
Theta rhythms and modulator identification. **(a)** LFP recordings across the sampled network (blue). Red lines show the theta frequency component (4-10 Hz) of the LFP signal for each presented site. Theta rhythms are clearly present across the sampled network (gray shaded boxes), often occurring for a second or longer with an intermittent dynamic. **(b)** Left: Microstimulation of the sender OFC does not evoke a receiver response at the M1 site due to disruptive neuromodulation of the sender-receiver communication by the M1 site identified as modulator. Right: Microstimulation of the sender OFC evokes a receiver response at the M1 site due to constructive neuromodulation of the sender-receiver communication by the M1 site identified as modulator. Center: Empirical distribution example of mean-log 𝛳 power for hit/miss receiver evoked responses at the M1 receiver site. **(c)** Schematic of the modulator decoding algorithm using baseline theta activity. TPR, true positive rate, FPR, false positive rate, P_FDR_, p-value corrected by false discovery rate analysis for multiple comparison.

Across the database of the 110 detected sender-receiver pairs in both monkeys, at least one significant modulator was identified in 36 sender-receiver pairs (36/110 = 33%). Overall, we identified 170 theta-modulators across the 110 recording sessions with a detected sender-receiver pair. **Table S2** lists the number of modulators found in each sender-receiver pair. We then defined positive modulators by asking whether increased modulator theta power was associated with increased sender-receiver communication and decreased modulator theta power was associated with decreased sender-receiver communication (**Methods**). Across the 170 modulator sites, 137 sites were positive modulators (80%). These results demonstrate that theta rhythms are associated with multiregional communication because they selectively predict stimulation responses and that increased/decreased modulator theta power predicted increased/decreased sender-receiver communication, respectively.

### Theta coherence and sender-receiver communication

We next investigated how theta coherence was associated with SR communication. Theta rhythms were selectively coherent across electrodes in different brain regions and at different times. An example of a simultaneous recording is shown in Fig 3a, with two exemplary events highlighted (shaded boxes). In one event, dPFC and M1 show theta coherence while in the other event CN and ACC show coherent theta. Examining all pairs of sites in a sample recording revealed that the grand average coherence was prominent in the theta frequency (4-10 Hz) and greater than that present in the beta frequency (15-25 Hz; Fig 3b). Thus, theta coherence is widespread supporting an important role in multiregional communication. Moment by moment, pairs of regions displayed coherent theta activity while others did not, demonstrating regional specificity.

**Figure 3.**
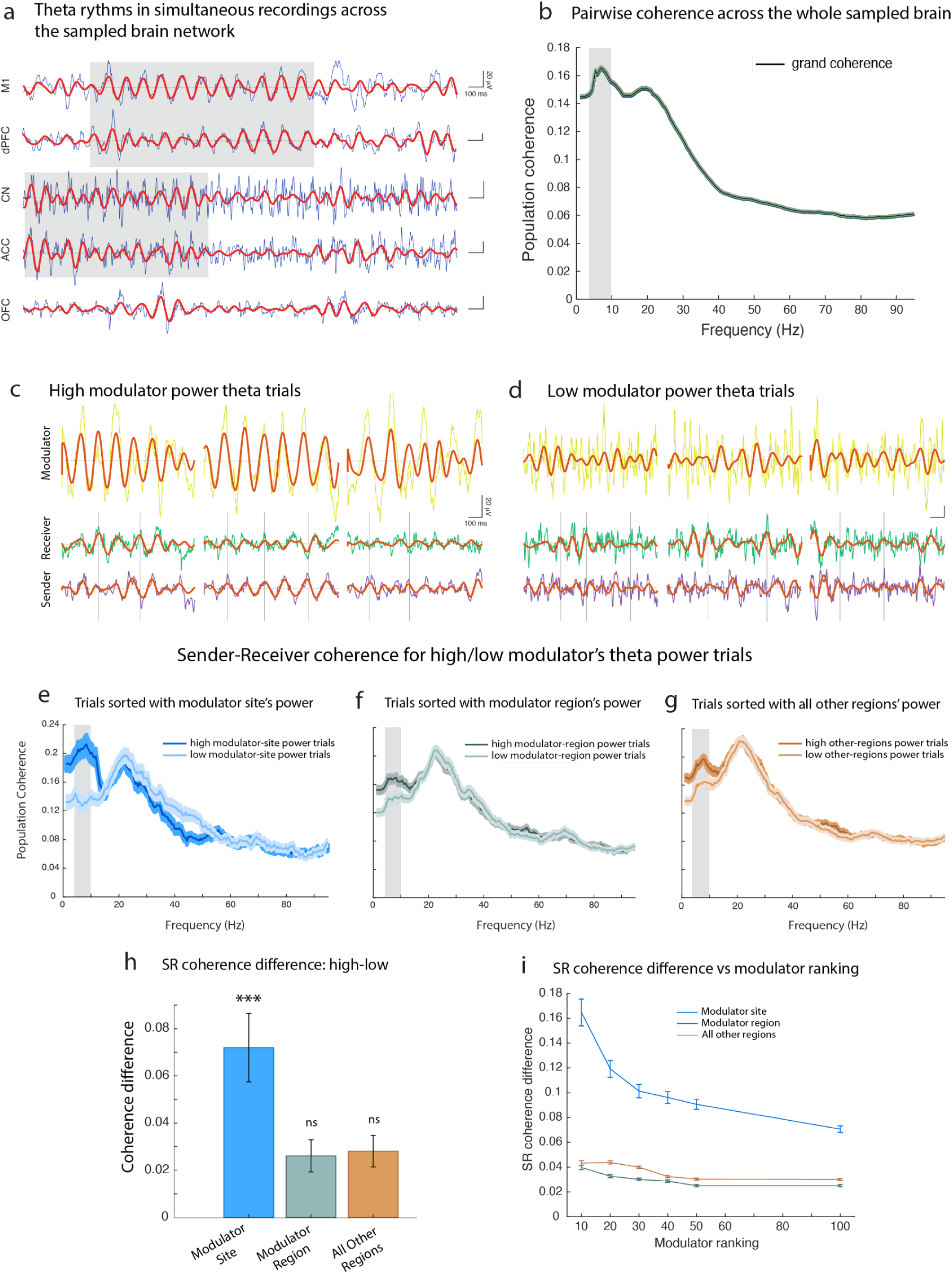
Theta rhythms and sender-receiver communication. **(a)** Examples of simultaneous LFP recordings in the sampled brain network (blue). The theta frequency component of the signal (4-10 Hz) is illustrated in red. Shaded gray boxes highlight theta-coherent events. **(b)** Pairwise coherence across all sampled recordings (error bars, SEM). Theta-frequency (4-10 Hz, grey shaded area) coherence is the most prominent coherence across the whole sampled brain network and it is greater than that present in the beta-frequency range (15-25 Hz). **(c)** Theta activity of the identified sender and receiver (bottom and center) during epochs with high theta power in the identified modulator site (modulator theta activity shown on top). **(d)** Theta activity of the identified receiver and sender (bottom and center) during epochs with low theta power in the identified modulator site (modulator theta activity shown on top). **(e)** Sender-receiver population coherence during time epochs with high and low theta power in the identified modulators (error bars ± SEM). SR theta-coherence was significantly greater in epochs with high modulator theta power compared to epochs with low modulator theta power (gray shaded bar, p = 0.03, permutation test). **(f)** Sender-receiver population coherence during time epochs with high and low theta power in the other sites within the modulator region (error bars ± SEM). SR theta-coherence is not significantly different in epochs with high modulator’s region theta power compared to low modulator’s region theta power (p=0.25, permutation test, modulator site(s) not included in the calculation). **(e)** Sender-receiver population coherence during epochs with high and low theta power at sites in other regions (error bars ± SEM). No significant difference was detected between SR theta-coherence in epochs with high theta power compared to epochs with low theta power in other brain regions (p=0.19, permutation test). **(h)** Sender-receiver coherence difference (high power trials minus low power trials coherence) in the theta range (gray shaded area) for the cases shown in panels (e) (***p=0.03, permutation test) and panels (f,g) (ns: p = 0.25, p = 0.19 respectively, permutation tests). **(i)** Sender-receiver theta-coherence difference vs modulator ranking. Consistent with the predictions, SR-coherence change with modulator power is a monotonic function of the modulators’ ranking and higher for modulators that better predict SR communication. Ranking by the theta power of other sites in the modulator region or other regions did not produce a similar change.

The results of theta power analysis (Fig 2) suggests that theta coherence may specifically reflect the modulation of SR communication. We therefore examined whether changes in SR theta coherence are predicted by theta power in modulator sites. Importantly, since theta coherence between the sender and the receiver and theta power in the modulator involve recordings in different electrodes, theta power in the modulator is not necessarily related to changes in SR theta coherence.

**Figures 3c and 3d** illustrate theta activity in example recordings with identified senders and receivers during time epochs with high and low theta power in the modulator, respectively. When modulator theta power was high, theta activity in the sender and receiver tended to fluctuate coherently but not when theta power was low (vertical lines). Across the population of recordings, SR theta coherence was significantly greater when modulator theta power was high compared to when modulator theta power was low (Fig 3e, p=0.03, permutation test, **Methods)**. This was not true for theta power at other sites in the modulator region (Fig 3f, p=0.25, permutation test) or at sites in other regions (Fig 3g, p=0.19, permutation test). Therefore, covariation of SR theta coherence with modulator theta power was specific to theta-frequency activity across the causally-identified sender-receiver-modulator (SRM) sites (Fig 3h).

Modulation of SR coherence may reflect modulation of communication between the sender and receiver. If so, changes in the power of modulators that more accurately predict changes in SR communication should be associated with changes in SR coherence. To measure the modulator’s predictive power we used the AUC obtained from the ROC analysis predicting receiver responses to sender stimulation from modulator activity immediately before stimulation. We then tested the prediction by ranking modulators and computing the change in SR coherence observed for high modulator power and low modulator power for each percentile (**Methods**). Consistent with the prediction, the change in SR coherence with modulator power was a monotonic function of the modulator ranking and greatest for modulators that most accurately predicted SR communication (Fig 3i). Importantly, similar changes in SR coherence were not observed when ranking either by theta power at other sites in the modulator region or by ranking by theta power at other sites in other regions. Consequently, changes in theta power at the strongest modulators, i.e. those more strongly predicting SR communication, best predict changes in SR coherence.

The tight relationship we observe between modulator theta power and SR theta coherence could be due to volume conduction and specifically associated with zero phase coherence^18^. If so, increasing modulator theta power should be associated with zero phase coherence. To address this concern, we compared SR theta coherence for trials with low modulator power and high modulator power (**Supp. Fig. S1**). When modulator power was low, we observed that SR theta coherence could be in-phase or out-of-phase and this did not significantly change when modulator power was high. Therefore, changes in SR coherence with modulator theta power could not be explained by volume conduction from the modulator site.

These results show that theta power in modulator sites is tightly linked to SR theta coherence. SR theta coherence tracks increases and decreases in neural excitability inferred from changes in theta power in the modulator, i.e. whether sender stimulation can generate a response in a receiver. Moreover, changes in SR theta coherence with modulator theta power are specific to the set of SRM sites identified by responses to electrical stimulation.

### Channel-modulation-by-coherence

The relationships between modulator theta power and SR communication support a conceptual framework in which modulation of excitability across the SR communication channel occurs due to changes in modulator theta power. When excitability is high, the sender communicates with the receiver (Fig 4a). When excitability is low, communication is suppressed (Fig 4b). What remains unclear, is the degree to which modulation of SR communication is specific to the phase of theta activity at the modulator site.

**Figure 4.**
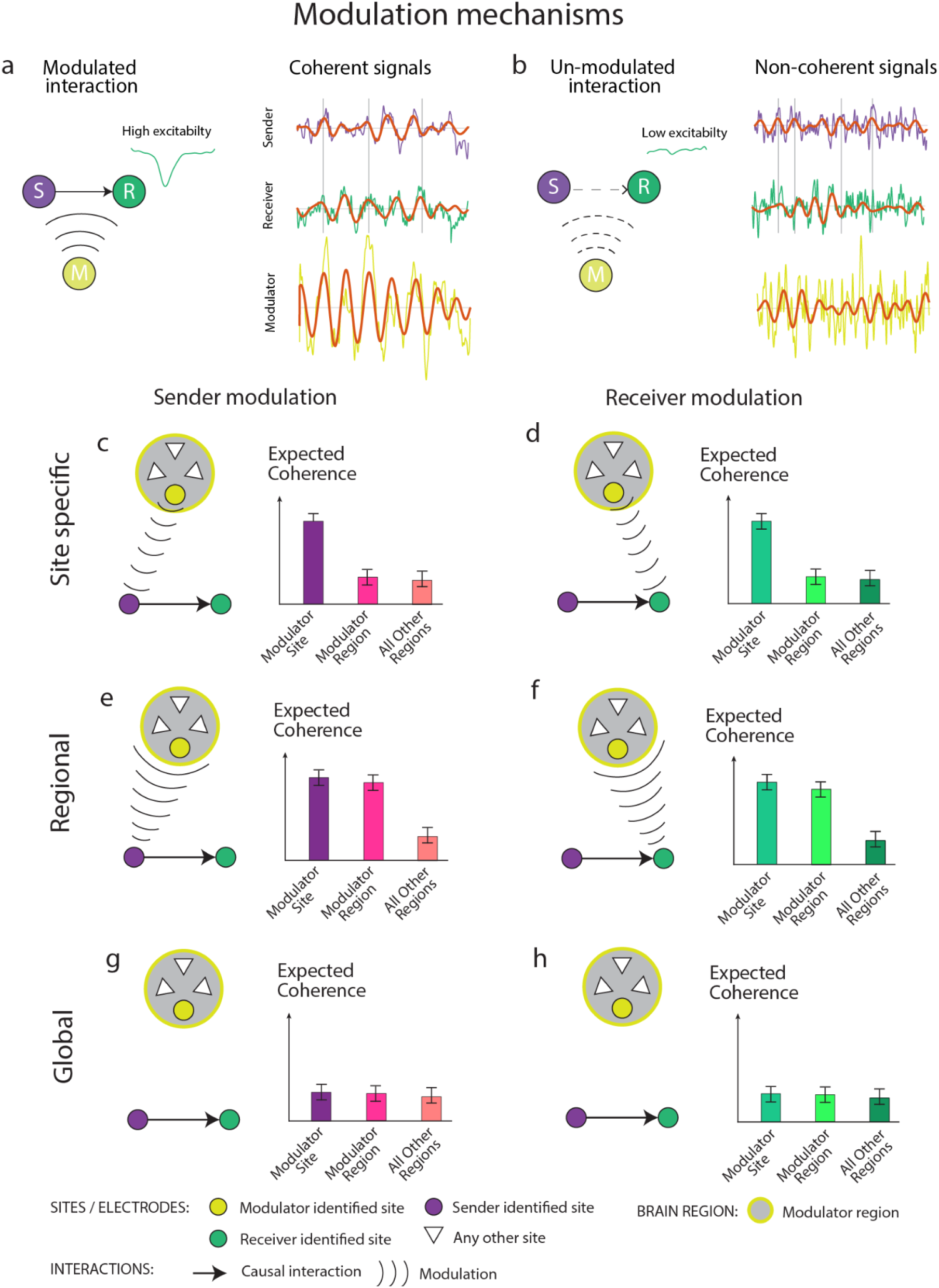
Modulation mechanisms. **(a)** Left: schematic sender-receiver modulated interaction. Constructive modulation favors high receiver’s excitability and, in turn, sender-receiver communication. Right: sender-receiver modulation occurs throughout coherent theta activity between the sender, receiver, and modulator sites. **(b) Left:** Schematic of sender-receiver modulated interaction. Destructive modulation interferes with sender-receiver interactions and favors low receiver’s excitability. Incoherent theta activity between sender, receiver, and modulator sites prevents sender-receiver communication. **(c) Left:** schematic of *site-specific hypothesis* of the sender modulation. The modulator theta phase couples with changes in neural excitability at the sender-site. **Right:** theta coherence between sender and modulator should be present in this case. **(d) Left:** schematic of *site-specific hypothesis* of the receiver modulation. **Right:** Similarly to (c), theta coherence between receiver and modulator should be present in this case. **(e) Left:** schematic of *region-specific hypothesis* of the sender modulation. The theta phase of all sites in the modulator region couples with changes in neural excitability at the sender-site. **Right:** theta coherence between sender-modulator-site and between sender-modulator-region should not differ in this case. **(f) Left:** schematic of *region-specific hypothesis* of the receiver modulation. The theta phase of all sites in the modulator region couples with changes in neural excitability at the receiver-site. **Right:** no difference between the receiver-modulator-site coherence and receiver-modulator-region coherence should be detected in this case. **(g-h)** schematic of *global hypothesis* of the sender and receiver respectively. According to this hypothesis, the coherence between the modulator-sender and modulator-receiver should not be different from coherence between the sender (receiver) and all other regions.

According to the *site-specific hypothesis*, if the modulator theta phase couples with changes in neural excitability at the sender site, then theta coherence between the modulator and sender sites should be present (Fig 4c). Similarly, if the modulator theta phase couples with changes in excitability at the receiver side of the channel then theta coherence should be present between the modulator and receiver sites (Fig 4d). Alternatively, theta phase across the modulator region couples with changes in sender and/or receiver excitability. According to this *regional hypothesis*, theta coherence with the sender site may not differ across sites in the modulator region (Fig 4e) and theta coherence with the receiver site may not differ across sites in the modulator region (Fig 4f). Finally, under the *global* hypothesis, theta coherence between the modulator and sender and between the modulator and receiver may not differ from the coherence with all other regions (Fig 4g**-h**).

To test these hypotheses, we computed the modulator-sender and modulator-receiver coherence across the population of all modulators associated with each SR pair (see **Methods**). We first consider modulator-sender coherence. Population average sender coherence with the modulator site, modulator region and all other regions are shown in Fig 5a and Fig 5b. *Sender-modulator-site* theta-frequency coherence magnitude was greater than the *sender-modulator-region* coherence (Fig 5b, p = 8*10^-4^, permutation test). Sender-modulator-region theta coherence was indistinguishable from theta coherence between the sender and all other regions (p= 0.06, permutation test, see Fig 5a, gray-band range and Fig 5b). Thus, channel modulation-related theta coherence is site-specific rather than region-specific (compare Fig 5b with Fig 4c). Coherence between the sender and other sites in the modulator region was no greater than between the sender and sites in any other region sampled in the network. Theta coherence with the sender sites was largely restricted to the modulators themselves (Fig. 5b). Figure 5c illustrates the site-specificity of the effects for example sender-modulator (SM) sites spanning CN, M1, and OFC.

**Figure 5.**
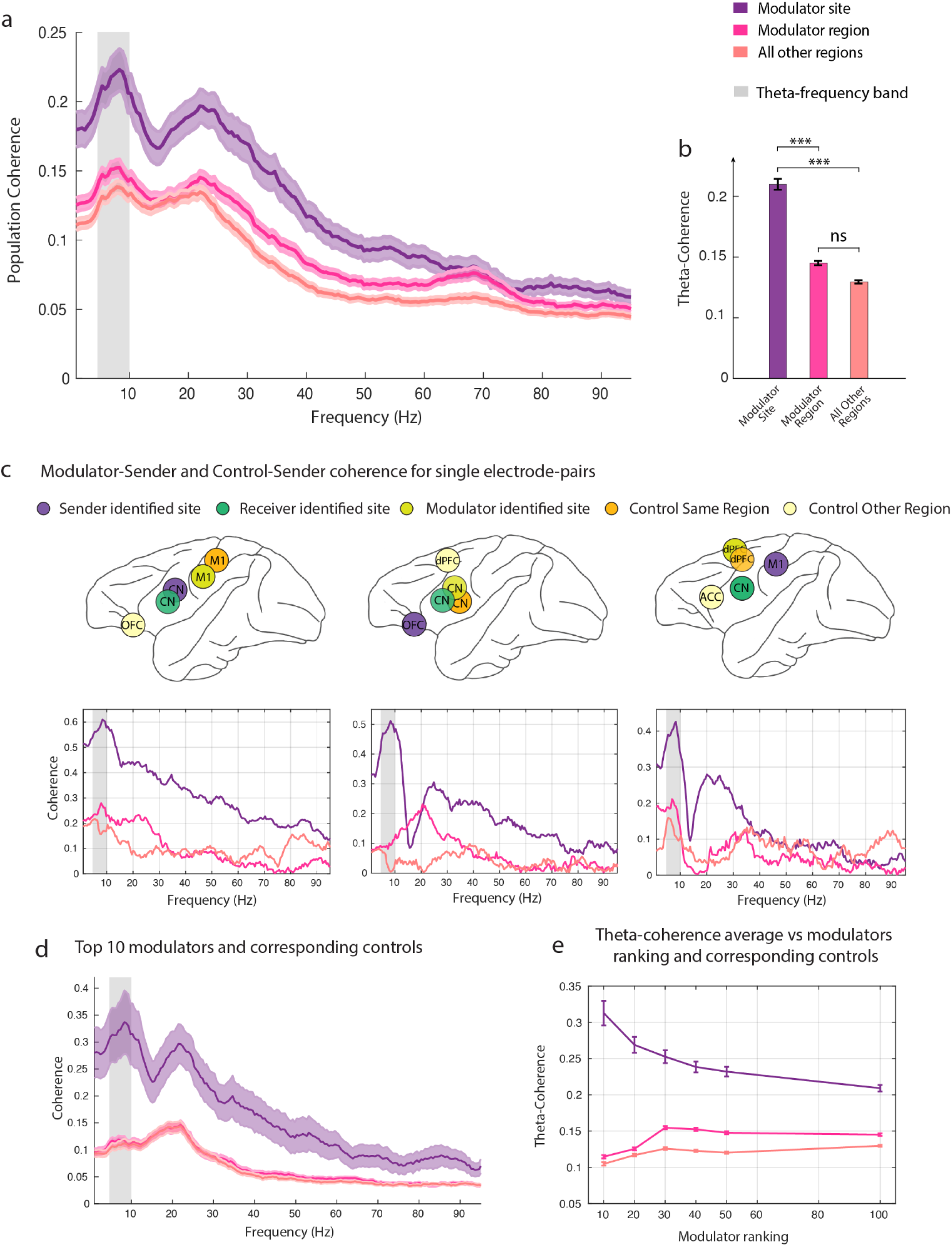
Sender modulation. **(a)** Population coherence between the sender-modulator-site, the sender-modulator-region, and the sender and all other regions, error bars ± SEM (gray shaded bar indicates the theta-frequency range 4-10 Hz). **(b)** Population theta-coherences in (a) (average results only in the theta-band) and permutation test results (**Methods**). SM coherence is significantly higher than sender-modulator-region and sender-other-regions coherence (***: p = 8×10^-4^ and p = 1×10^-4^, respectively). Sender-modulator-region coherence is not distinguishable from sender-other-regions coherence (ns: p = 0.06). All statistical significance is assessed via permutation tests. **(c) Bottom**: Coherence examples for electrode in Modulator-Site and Sender, electrode in Modulator-Region and Sender, electrode in other regions and Sender. **Top**: illustration of electrodes location/brain region. **(d)** Population coherence between the sender-modulator site, the sender-modulator-region, and the sender and all other regions (as in panel a) but including only the top 10 ranked modulators (**Methods**). **(e)** Theta-coherence (average in [4-10] Hz range) as a function of the modulator ranking for the sender-modulator-site, the sender-modulator-region and the sender and all other regions. SM theta coherence monotonically decreases as a function of the modulator ranking and it is significantly higher than modulator region and all-other-region alternatives.

We next tested the association between modulator predictive power and coherence with the sender. To measure the modulator’s predictive power we used the AUC obtained from the ROC analysis predicting receiver responses to sender stimulation from modulator activity immediately before stimulation. We then ranked modulators and averaged the sender coherence with the modulator site for each percentile (**Methods**). Consistent with the site-specific hypothesis, when including only the top 10% of ranked modulators, the SM-site coherence is much greater than when including all the modulators, especially around the theta frequency (Fig 5d). There was no difference between the modulator region coherence and all-other-region coherence with the sender, rejecting both regional and global hypotheses for sender modulation. Across the population, SM theta-coherence monotonically decreased as a function of the modulator ranking and was significantly greater than modulator region and all-other-region alternatives (Fig 5e). Consequently, the strongest modulators, i.e. those more strongly predicting SR communication, are most highly coherent with the sender.

We then performed the equivalent analyses for receiver sites to assess whether Receiver-modulator-site (RM) theta coherence may also reflect channel modulation. As for sender sites, RM coherence was significantly higher for the modulator site than the modulator region (Fig 6a**-c**, p=5*10^-4^, permutation test). Theta coherence after ranking modulators reinforced the result, showing that modulation of the receiver is *site-specific* rather than *region-specific (*Fig 6d**,e**). Taken together, these results demonstrate that theta-coherence could modulate SR communication in a site-specific manner.

**Figure 6.**
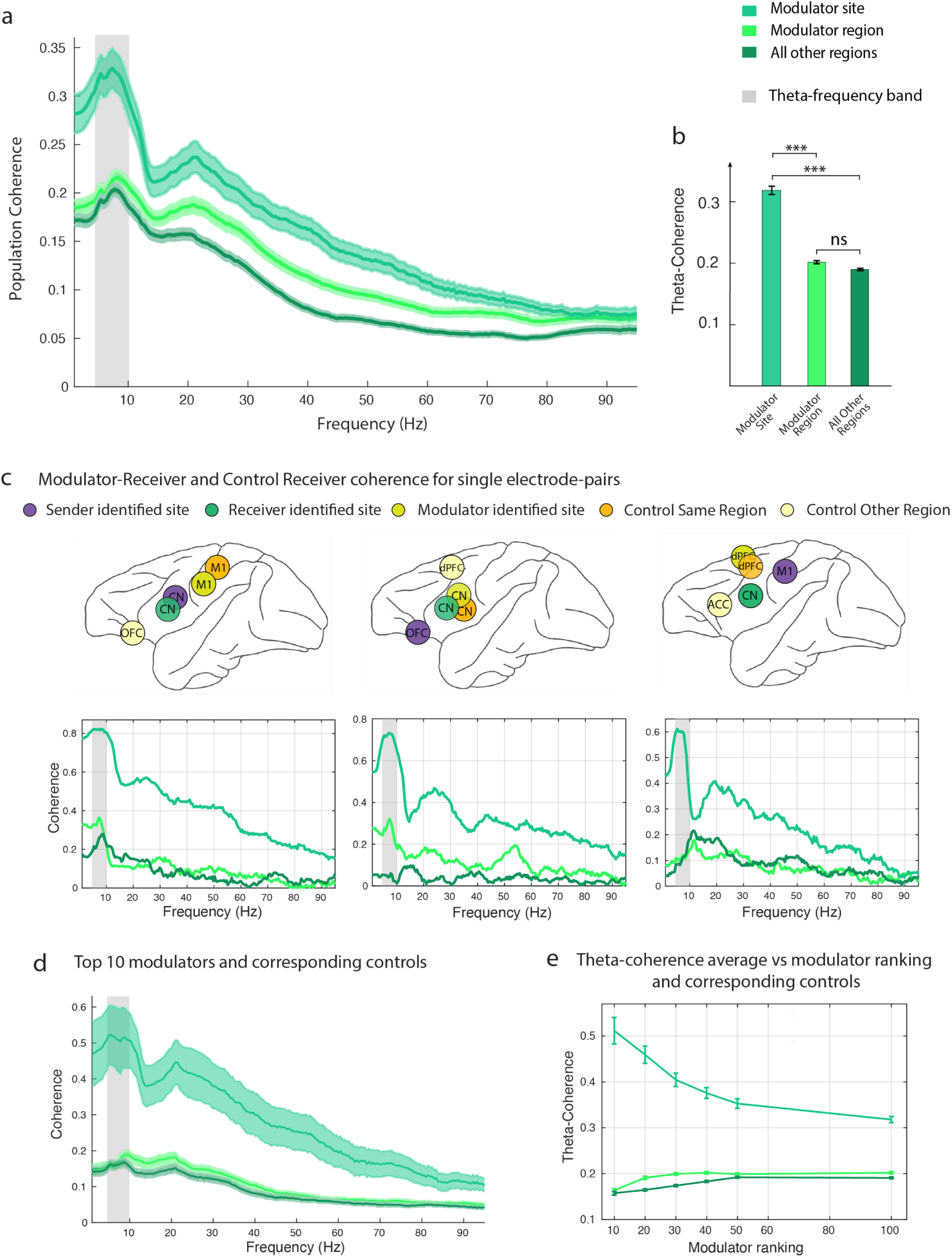
Receiver modulation. **(a)** Population coherence between the receiver-modulator-site, the receiver-modulator-region, and the receiver and all other regions, error bars ± SEM (gray shaded bar indicates the theta-frequency range 4-10 Hz). **(b)** Population theta-coherences in (a) (average results only in the theta-band) and permutation test results. Population theta-coherence comparison: RM coherence is significantly higher than receiver-modulator-region and receiver-other-regions coherence (***: p = 5*10^-4^ and p = 10^-4^, respectively). Receiver-modulator-region coherence is not distinguishable from receiver-other-regions coherence (n.s.: p = 0.21). All statistical significance is assessed via permutation test (**Methods**). **(c) Bottom**: Coherence examples for electrode in Modulator-Site and Receiver, electrode in Modulator-Region and Receiver, electrode in other regions and Receiver. **Top**: illustration of electrodes location/brain region. **(d)** Population coherence between the receiver-modulator site, the receiver-modulator-region, and the receiver and all other regions (as in panel a) but including only the top 10 ranked modulators (**Methods**). **(e)** Theta-coherence (average in [4-10] Hz range) as a function of the modulator ranking for the receiver-modulator-site, the receiver-modulator-region and the receiver and all other regions. RM theta coherence monotonically decreases as a function of the modulator ranking and it is significantly higher than modulator region and all-other-region alternatives.

Directly comparing SM coherence and RM coherence, we observe that the coherence magnitude is greatest in the theta-frequency band (∼8 Hz peak, Fig. 7a). RM theta coherence magnitude is ∼52% stronger than SM theta-coherence. Figure 7b summarizes the key result that modulator theta activity is coherent with receiver activity more than with sender activity. The overall pattern of results suggests that changes in modulator theta power that can modulate SR communication are coherent with the receiver site more than the sender site.

**Figure 7.**
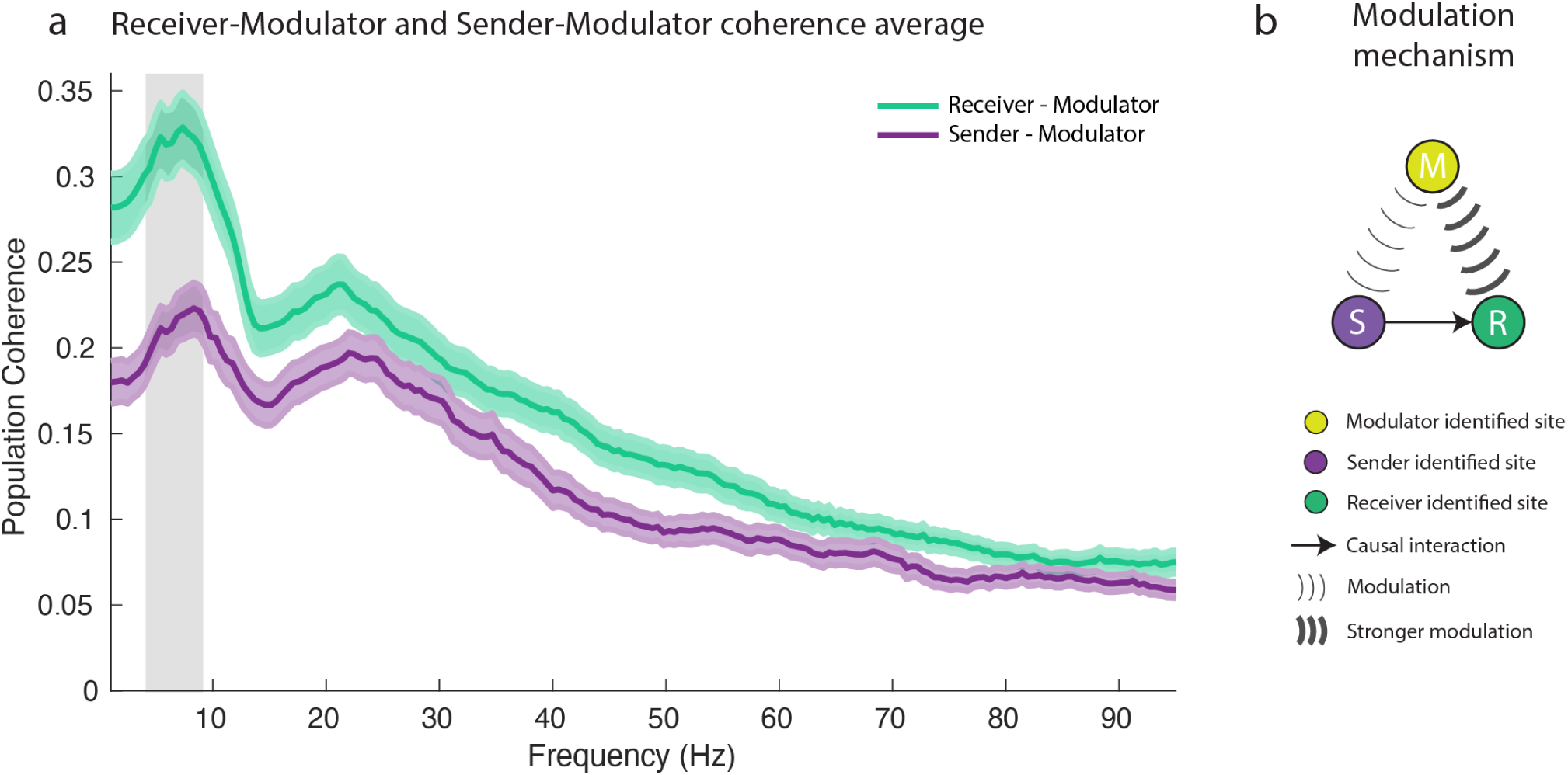
Summary of the modulation mechanism: **(a)** Direct comparison of population RM- and SM-coherence. RM theta coherence magnitude is ∼52% stronger than SM theta-coherence (gray shaded region). **(b)** Modulator mechanism: Modulator theta activity is more coherent with the receiver-site activity than with the sender-site activity. Changes in the modulator theta power that can modulate sender-receiver activity are coherent with the receiver site more than with the sender-site.

### Distributed modulator networks

The prominence of theta coherence across SRM motifs suggests that theta coherence could coordinate multiregional communication across multiple modulator regions that form a modulator network. Some of the SR pairs that we identify via the stimulation experiment are modulated by more than one site (**Table S1**, Fig. 8a for monkey M). We identified as many as 18 modulators for a single SR pair. We used this sample to investigate the pairwise theta-coherence structure among the modulators and to test for the existence of a modulator network associated with a particular SR communication channel.

**Figure 8.**
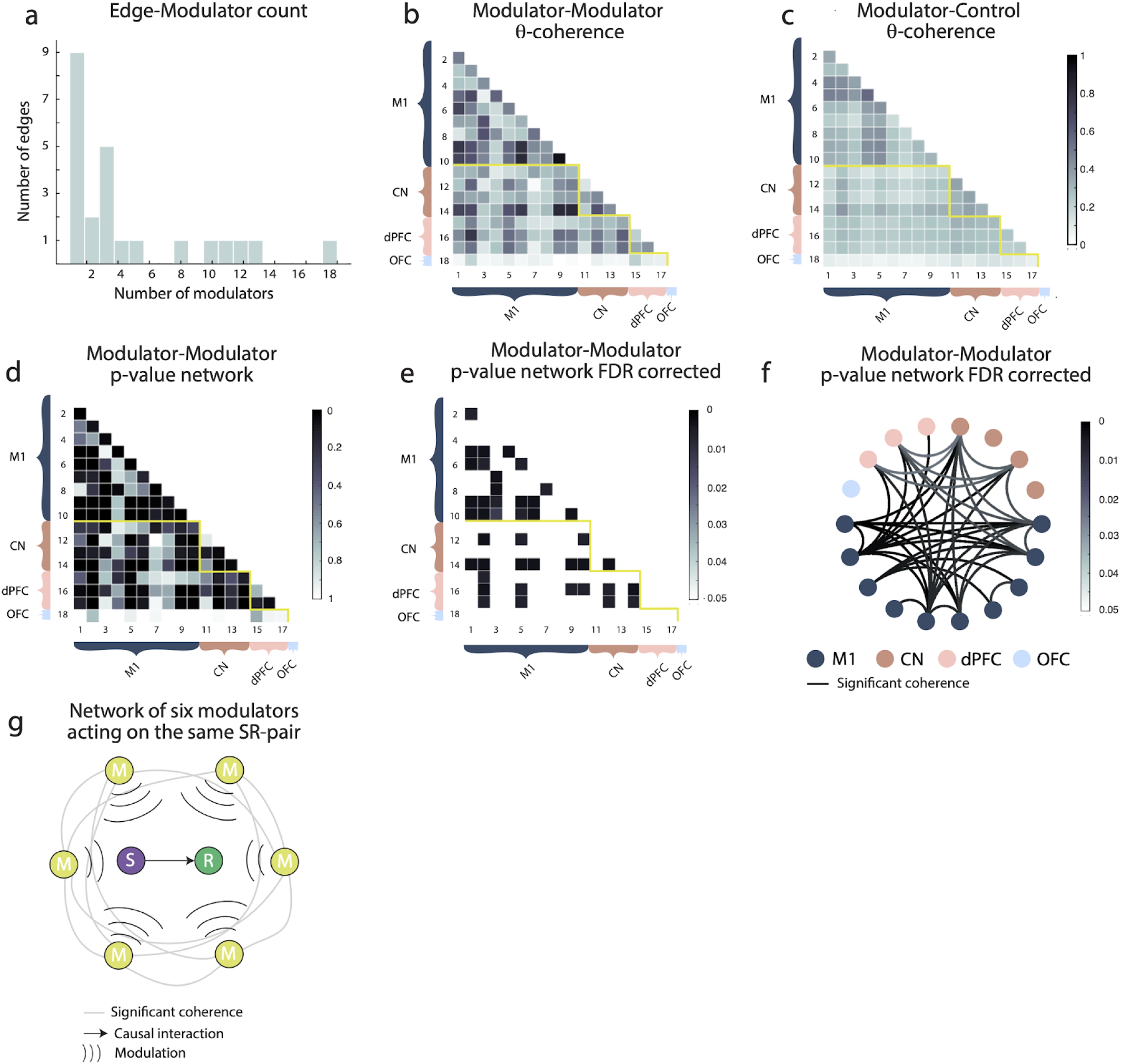
Network of modulators. **(a)** Empirical distribution of the number of sender-receiver pairs with a given modulator count. **(b)** Modulator-modulator pairwise theta coherence for a session with the largest number of modulators (18 sites). Electrodes are organized by brain region, as illustrated in the panel. **(c)** Modulator-Control pairwise theta-coherence for the same session considered in (b) used to create a null-distribution for the modulator-modulator theta coherence. **(d)** Statistical significance (p-value) of the modulator-modulator theta-coherence shown in panel **(a)**. **(e)** Same as in (d) after FDR correction for multiple comparison (α = 0. 05), values represent P_FDR_ . **(f)** Network representation of modulator-modulator statistical significant theta-coherence (network representation of the matrix shown in **(e)**: the existence of a link represents a statistical significant theta-coherence among a pair of modulators). **(g)** Illustration of a theta-coherent network of 6 modulators modulating a causal sender-receiver pair. All results for Monkey M. See **Supp Fig. S2** for Monkey A.

To characterize the modulator network associated with the SR channel, we calculated the pairwise modulator-modulator (MM) theta-coherence across all modulator sites (Fig. 8b, **Methods**). Figure 8b presents the results for modulator sites grouped anatomically: electrodes in the same brain region are listed contiguously with yellow lines marking the separation between two different brain regions. We then tested whether the MM coherence reflects an underlying modulator network. To do so, we computed the modulator coherence that controls for the regions of interest, termed the modulator-control coherence, defined as M_ij_C for a pair of modulators M_i_ and M_j_ (Fig 8c**, Methods**). We then used the null distribution to estimate statistical significance for each M_i_M_j_ coherence (Fig. 8d) and controlled for multiple comparisons (Fig 8e, **Methods,** P_FDR_<0.05). Modulators associated with a SR channel are significantly coherent among themselves. Interestingly, examining the anatomical distribution of the modulator network revealed that modulator coherence is significant within and between regions (Fig 8f).

These results indicate that, theta-coherence associated with modulation of multiregional communication involves both every modulator and the SR-pair and, concurrently, the modulator-modulator multiregional network (Fig. 8g provides a schematic for such interpretation). Similar results for Monkey A are presented in **Supp. Fig. S2**.

## Discussion

Here, we test the link between theta rhythms, multiregional theta coherence and multiregional communication. Using simultaneous multiregional recordings and electrical microstimulation across the primate mesolimbic mood network, we show that sender and receiver sites whose communication is identified causally also display strong theta-frequency coherence. We discovered that pairs of causally-identified sender and receiver sites exhibited more coherent theta rhythms than other nearby sites. We also show that moment-by-moment changes in sender-receiver communication can be identified using changes in theta-band LFP power at modulator sites. Specifically, we identify modulator brain sites whose theta power predicts variability in the receiver response to sender stimulation, and that modulator-sender-receiver subnetworks display distributed theta coherence. These results provide important causal evidence that coherent theta-activity could play a direct role in the control and modulation of multiregional communication.

The finding that neural modulation of sender-receiver causal interaction, which we term channel modulation, could be conveyed through theta-coherence between the modulator sites and the sender-receiver pair is particularly important. Modulator sites with the highest predictive accuracy (modulator ranking) of the SR-interaction are also those more highly theta-coherent with both the sender and the receiver site and that, further, theta-coherence monotonically increases with the modulator ranking. Thus, channel modulation is strongly linked to theta-coherence and stronger modulation implies greater coherence and vice-versa.

Assessing the extent to which theta coherence between the modulator and the sender regions reflects diffuse indirect interactions shared between sites across each regions or, alternatively, reflects more direct properties specific to the modulator sites themselves has important implications for understanding the nature of theta coherence and the role of modulator sites in multiregional communication. We find that dynamic theta-frequency-related modulation of multiregional communication is more site-specific than region-specific: High theta-coherence is not present diffusely from one region to another and it rather varies from site-to-site. Site-specificity suggests that specific populations of neurons in the modulator regions may modulate sender-receiver communication. These populations may be in the specific layers from which modulator projections arise. Site-specificity may also reflect patchy distributions of connections from groups of neurons in the modulators and well the presence of locally connected groups of neurons in the senders/receivers.

### Theta rhythms and multiregional communication

Theta activity in the brain is widespread across disparate brain regions during high-level cognitive processes^19^. A strong connection exists between theta and memory encoding and retrieval in the frontal lobe, working memory, and top-down control in the frontal lobe ^20–23^. Multielectrode intracranial EEG (iEEG) demonstrates theta oscillations occur broadly across the human cortex and in different behavioral states^21,23–27^. Theta oscillations have also been shown to engage multiple brain regions while playing a role in human spatial navigation. Magnetoencephalography reveals that theta oscillations were increased in the parahippocampal gyrus and posterior cingulate cortex (PCC) during navigation^28^. Theta power in the PCC has also been positively correlated with navigation performance, with oscillations not exclusively related to movement or speed^29^.

In non-human primates, theta oscillatory activity has been implicated in multiregional communication across a variety of different behavioral tasks^30,31^ and features of theta rhythms in area 9 (the medial prefrontal cortex) and area 32 (the rostral ACC) are associated with “executive attention” including self-control, internal timing, and assessment of reward^32^. Studies in humans and non-human primates have also linked attention-related perceptual changes to intrinsic dynamics of theta rhythms^1,33–35^. In each case, theta rhythms have been associated with multiregional communication.

Our findings use simultaneous multiregional recordings and microstimulation to define specific roles for brain sites as senders, receivers, and modulators and suggest that changes in theta power and coherence can be interpreted in terms of changes in multiregional communication. These considerations suggest modulator-sender and modulator-receiver coherence can serve as a marker of when sender-receiver communication can occur. We find that modulator sites are theta-coherent with both the sender and the receiver: their theta-coherence is significantly higher than control coherence. This supports the importance of coordinated activity between the modulator and the sender-receiver pair and further shows that interregional modulation involves both sites in the SR-pair. Furthermore, we find that theta-coherence is significantly much higher between the modulator-receiver than the modulator-sender pair. Therefore, while modulation involves both sender and receiver, coordinated theta-rhythms are stronger between the modulator-receiver than between the modulator-sender pair.

### Coherent theta rhythms play a pivotal role in brain communication and neuromodulation

This work sheds light on the integral roles of theta rhythms and coherence in brain communication and neuromodulation, highlighting their fundamental significance within sender-receiver-modulator (SRM) triads. Our results demonstrated that the theta power of modulator sites could effectively predict the receiver’s response to sender stimulation. High modulator theta power was associated with an increased receiver response, suggesting a pivotal role for modulators in controlling the flow of information in sender-receiver neural circuits. In addition, high modulator theta power corresponded to high SR theta coherence, indicating that theta rhythms are implicated in regulating the efficiency of SRM communication channels.

Importantly, we found that the most influential modulators, i.e. those with the highest rankings, were associated with larger theta coherence within the SRM triads. This suggests that the strength of theta coherence is intimately linked with the potency of the modulation process, reinforcing the pivotal role of theta rhythms in regulating multiregional communication. Our work complements recent results showing that short-burst, 2 s, 100 or 200 Hz, tetanic electrical microstimulation at modulator sites can alter the sender-receiver channel of communication by disrupting receiver’s excitability^36,37^. This offers direct evidence of a causal relationship between the modulator and the sender-receiver pair. This result, together with our findings, suggests that high modulator’s theta power causally facilitates SR communication which, in turns, happens through SR theta synchronization. Furthermore, we showed that the SRM triads are significantly theta-coherent compared to other brain sites which, combined with the SRM causal relationships, illustrate a pivotal role for coherent theta rhythms in the dynamic control of multiregional communication and neuromodulation.

### Manipulation of theta rhythms through deep brain stimulation

The role of theta rhythms in cognitive processes means disordered theta activity has been implicated in neuropsychiatric disorders including anxiety, obsessive compulsive disorder (OCD), and depression^38,39^. In anxiety disorders, frontal-midline theta signals reflecting midcingulate cortex activity are moderated by anxiety in both resting state and task conditions^40^ with anxious individuals showing greater theta activity. In OCD, elevated theta power has been observed in frontal and central regions, suggesting hyperactivity in the cortico-striato-thalamo-cortical loop, a key circuit implicated in the pathophysiology of OCD^41^. Altered theta activity, particularly in the ACC and in the superior temporal gyrus has been consistently reported in depressed patients^42,43^. In rostral ACC, altered theta activity can predict pharmacological and behavioral treatment outcome ^42^ and may serve as a biomarker^44^.

Deep brain stimulation (DBS) is an important approach to treating otherwise resistant forms of brain disorders in epilepsy^45^, treatment-resistant depression^46^, OCD^47^, and chronic pain^48^. DBS is also used to alter brain rhythms including theta-rhythms, in order to restore or enhance cognitive functions^49^. A more precise understanding of how theta rhythms operate across modulatory brain networks may help design better DBS treatments aimed at regulating theta oscillations and it may lead to the development of more effective therapies for neurological and psychiatric disorders. This would allow for more targeted modulation of specific communication channels, potentially ameliorating symptoms of disorders characterized by dysregulated brain communication. Consequently, our work not only furthers our understanding of the functional significance of theta rhythms in brain communication but also has the potential to translate into clinically meaningful applications.

## METHODS

### SUBJECT DETAILS

Two male rhesus macaques (*Macaca mulatta*) participated in the study (Monkey M, 8.4 Kg and Monkey A, 7 kg). All surgical procedures were performed in compliance with the National Institute of Health Guide for Care and Use of Laboratory Animals and further approved by the New York University Institutional Animal Care and Use Committee.

### EXPERIMENT DETAILS

#### Microdrive stimulation-recording system

The microdrive system used for this study was developed and customized for intracortical microstimulation (ICMS) and recording of the cortico-subcortical mood processing network^37^. The screw-driven actuation mechanism controlled the position of 220 microelectrodes bidirectionally and independently (1.5 mm spacing) along a single axis with a range up to 32 mm (Monkey M) and 40 mm (Monkey A) with 125-μm pitch ^50^. We distributed two types of microelectrode in the microdrive for Monkey M: extracellular recording and ICMS was done through 160 platinum/iridium (Pt/Ir) electrodes (MicroProbes,Gaithersburg, MD) with impedance 0.1–0.5 Ω, 60 additional tungsten electrodes (Alpha Omega, Israel) with impedance 0.8–1.2 Ω were used for extracellular recording only. For Monkey A, we loaded 220 Pt/Ir electrodes (MicroProbes,Gaithersburg, MD) with impedance 0.5 Ω for extracellular recording and ICMS. Electrode impedance was measured at 1 kHz (Bak Electronics, Umatilla, FL). The shank diameter for the tungsten electrode was 125 mm (total diameter 250 mm with glass insulation). The shank diameter of the Pt/Ir electrode was 225 mm (total diameter 304 mm with parylene C and polyimide insulation).

#### Experimental preparation

Surgical and experimental methods are described in detail in previous work^37^. A craniotomy and dura thinning on the targeted brain regions was performed over the left hemisphere in Monkey M and over the right hemisphere for Monkey A. A customized large-scale recording chamber (Gray Matter Research, Bozeman, MT) was implanted and fitted to the skull’s surface using MR-guided stereotaxic surgical techniques (Brainsight, Rogue Research, Montreal, QC). The chamber was aligned and registered within 1 mm of the target coordinates (nominally 0.4 mm) and sealed to the skull surface via C&B-METABOND (Parknell Inc., Edgewood, NY) and dental acrylic. The microdrive was mounted into the chamber and sealed with compressed gaskets and room-temperature-vulcanizing (RTV) sealant (734 flowable sealant, Dow Corning, Midland, MI). To target each brain region, electrodes were registered through anatomical magnetic resonance images (MRIs) and magnetic resonance angiograms (MRAs). In order to limit routes for infection, the chamber and microdrive were customized to each animal’s anatomy. To limit damage to the vasculature, midline, and ventricles when lowering electrodes, only a subset of electrodes along direction considered safe were advanced. Structural images were co-registered to the MNI Paxinos labels^51^. We successfully studied activity from 165 electrodes (50 tungsten and 115 Pt/Ir) in Monkey M for over 24 months and 208 Pt/Ir electrodes in Monkey A for over 12 months.

#### Neurophysiology

The monkeys were awake, head-restrained and seated in a primate chair placed in an unlit electromagnetically shielded sound-attenuated booth (ETS Lindgren). Neural recordings were referenced to a ground screw in contact with the dura mater over the left posterior parietal lobe (Monkey M) or left occipital lobe (Monkey A). Neural signals from all channels were simultaneously amplified and digitized at 30 kHz with 16 bits of resolution with the lowest significant bit equal to 1 uV (NSpike, Harvard Instrumentation Lab; unit gain headstage, Blackrock Microsystems), and continuously streamed to disk during the experiment.

Microstimulation was applied using a bipolar configuration, made by sending a biphasic charge-balanced square wave pulse via a pair of Pt/Ir microelectrodes with the same pulse amplitude, pulse width, and interphase interval, but opposite polarity (e.g., cathode-lead for electrode 1 and anode-lead for electrode 2) (Cerestim R96, Blackrock Microsystems). The pulse width used was 100 ms per phase and interphase interval was chosen to be 53 ms for all stimulation sessions. The Pt/Ir microelectrodes had a typical tip geometric surface area of 223 ± 37 mm^2^. For example, a single pulse, with amplitude of 40 mA and width of 100 ms per phase (4 nC/phase), could yield a charge density of approximately 1800 mC/cm^2^. We simultaneously recorded neural signals from all electrodes while stimulating a certain pair of electrodes. No seizure activity due to stimulation was detected during experiments.

### Data analysis

#### Pulse-triggered evoked potentials

The LFP activity was obtained by low-pass filtering a raw signal recorded at 400 Hz using a multitaper filter with 0.025 s duration in time and bandwidth frequency of 400 Hz, center frequency of 0 Hz, and then decimating the signal 1 kHz from 30 kHz. In order to remove artifacts, we removed events which exceed 10 standard deviations from the mean across the stimulation event pool. Noisy and bad channels were also confirmed and removed by visual inspection. We computed pulse-triggered evoked potentials by averaging bipolar re-referenced LFP signals aligned to the onset time of each single pulse and then z-scored these signals using the standard deviation (SD) of the baseline (−30 to −5 ms). The SD of each electrode was computed separately using all time points in the baseline window of the pulse-triggered evoked potential.

#### Detection of sender-receiver causal interaction

The detection of the receiver’s evoked response to the single-pulse ICMS of the sender was quantified via the stimAccLLR algorithm^36,37^. The approach checks when selectivity in the LFP activity emerges by comparing the null to the alternative hypotheses, i.e., the latency of a stimulation response. Latency from a single microstimulation pulse was determined by measuring the time at which Pulse Event reached a detection threshold, which is selected based on a trade-off between speed (latency) and accuracy (probability of correct classification). The threshold is further used to determine the response to latencies by maximizing the difference between probability of ‘Hit’ and ‘Miss’ trials.

The stimAccLLR method is based on a probabilistic model of the LFP activity which tests for two alternative hypothesis, these are: H_1_, which represents the LFP activity in the post-stimulus epoch (Pulse Event), and H_0_, which represents the LFP activity in the pre-stimulus epoch (Pulse Null). We performed stimAccLLR on all stimulation responses to single microstimulation pulses.

The stimAccLLR method models LFP activity as independent observations from an underlying Gaussian distribution. The signal 𝑥(𝑡) at time is formalized as a mean function µ(𝑡) plus additive Gaussian noise ε(𝑡).

We estimated the mean response µ(𝑡) by low-pass filtering the raw LFP activity at 100 Hz

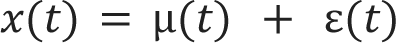

The likelihood of observing data 𝑥(𝑡) generated from such model is given by the Gaussian:

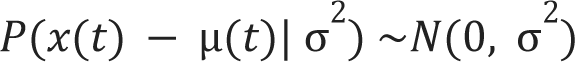

where ^2^ indicate the variance of the distribution. By using the above likelihood for two time-varying Gaussian LFP models we can calculate the likelihood ratio 𝐿𝑅(𝑡):

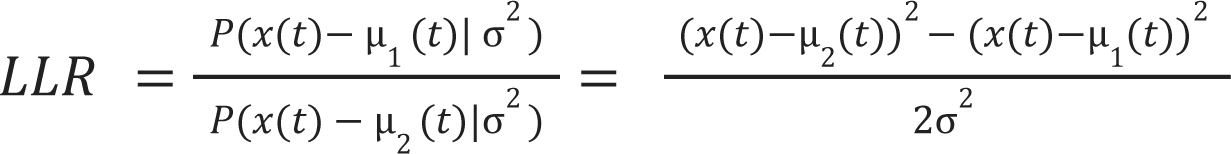

where we assumed that both models have the same variance σ^2^. When we estimated the 𝐿𝑅(𝑡) across events/trials we used leave-one out procedure. The accumulated log-likelihood ratio is then calculated by summing the log-likelihoods over time

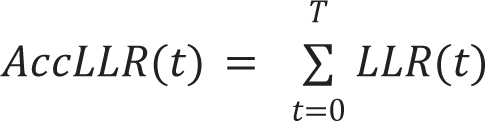

Signal selectivity was performed through a receiver-operating characteristic (ROC) analysis on the AccLLR values every millisecond, to discriminate between H_1_ (Pulse Event) and H_0_ (Pulse Null). An ideal observer analysis was used to measure the overall signal selectivity (OSS). We defined OSS as the choice probability from the ROC analysis at the end of an accumulation interval up to 100 ms. To correct for multiple comparisons, we used a false discovery rate (FDR) procedure ^52^ with an alpha threshold set at 0.01. This procedure yielded 110 significant sender-receiver pairs (edge samples, P_FDR_ < 0.01)^37^.

### Decoding sender-receiver causal interaction

Modulators were identified by decoding the receiver response to single-pulse stimulation of the sender that was detected using the StimAccLLR algorithm. For each identified sender-receiver pair and for each trial, the receiver’s response is classified either as a ‘hit’ (there is a significant response) or as a ‘miss’ (no significant response) according to the stimAccLLR result. We analyzed all simultaneously-recorded electrodes to decode the presence of the receiver response (hit/miss) from baseline activity (500 ms before pulse onset). We estimated LFP theta-power ([4 - 10] Hz) during the baseline epoch using a multitaper method with 500 ms sliding window and ±2 Hz smoothing. To reduce the presence of artifacts in the analysis, events that exceed 5 standard deviations from the mean across the simulation event pools were removed. For each electrode, we built the mean log θ-power histogram (Fig. 2b**-center**). We then perform a ROC analysis to test whether the LFP θ-power of each ‘hit’ trial can be distinguished from ‘miss’ trials (Fig. 2c). For each electrode, we compute the 𝑝-value of the area under the curve (AUC). To control for multiple comparisons across all the simultaneously tested sites, we performed the FDR procedure on all the tested sites for a given sender-receiver detected pair. A site is identified as a modulator if P_FDR_ of that site is P_FDR_ < 0.05 (Fig. 2c). Of all the 110 sender-receiver detected pairs, we tested 6189 sites as modulator candidates and we identified 170 significant theta-modulators (i.e. able to predict the receiver’s response throughout their baseline’s theta-power). The number of modulators for each sender-receiver pair is presented in **Table S2** and the modulator’s brain region list is presented in **Table S3**.

### Identification of positive modulators

Positive modulators are identified as those sites for which increased modulator theta power is associated with increased sender-receiver communication (i.e. more hits than misses) and decreased modulator theta power is associated with decreased sender-receiver communication (i.e. more misses than hits). To identify these modulators, for all the trials in each session (i.e. for each SR-pair), we compute the theta-power distribution of each modulator within the trial’s baseline activity (500 ms before sender stimulation). We define low-power trials those in the lower 1/3 of the power distribution, while high-power trials are defined as those in the higher 1/3 of the power distribution. We then construct a confusion matrix C whose entries are C_HH_ (count of hit trials for the receiver which are also modulator high-power trials); C_HL_ (count of hit trials for the receiver which are also modulator low-power trials); C_MH_ (count of miss trials for the receiver which are also modulator high-power trials); C_ML_ (count of miss trials for the receiver which are also modulator low-power trials). We define a positive modulator as a site for which C_HH_ + C_ML_ > C_HL_ + C_MH_ (diagonal sum > off-diagonal sum of the confusion matrix). Across the 170 modulator sites, 137 sites were positive modulators (80%).

### Sender-Receiver coherence and modulator theta power

To assess the statistical significance of the difference of the SR population coherences presented in Fig. 3e**-h** we performed a non-parametric statistical analysis. The test-statistics is defined as the average difference between the SR-theta coherence for high-power trials minus the same coherence for low-power trials (Fig. 3e**-h**, gray shaded area). A null distribution for the SR-coherence is built by randomly shuffling the high-power trials with the low-power trials and computing a new pseudo-SR-coherence difference for the shuffled data. For each session, we generate 1000 pseudo-coherence differences (1000 shuffles of high/low-power trials). The total number of sessions across monkeys amounts to 36, therefore the null distribution consists of 36000 pseudo-SR-coherence values. When we compute the modulator region theta power, the modulators present in the region are excluded from the calculation since they are only included in the modulator-site calculation.

### Sender-Receiver coherence difference for high/low modulator’s theta power trials as a function of the modulator ranking

We established the ranking of the modulator based on the Area Under the Curve (AUC) derived from the Receiver Operating Characteristic (ROC) analysis. This analysis predicts the response of the receiver to the sender’s stimulation based on the theta power activity of the modulator before the stimulation (using a 500 ms baseline period).

To calculate the difference in SR-theta coherence for high/low power trials at the modulator site (as shown by the blue line in Fig. 3i), we used a similar method to Fig. 3e. However, in this case, we only considered high/low power trials at the modulator site for a given fraction of the ranked modulators. The gray-green line in Fig. 3i represents the SR-theta coherence difference for high/low power trials across all electrodes in the modulator region(s). Here, we considered the same fraction of modulator-sites as in the case of the blue line.

Lastly, the brown line in Fig. 3i shows the SR-theta coherence difference for other regions. For this calculation, we included all the sites in other regions corresponding to the same fraction of modulator-sites considered for the blue line.

### Sender-Modulator coherence

For each sender-site S_j_ we compute the *sender-modulator-site* (S_j_ M_i_) coherence magnitude as a function of the frequency f:

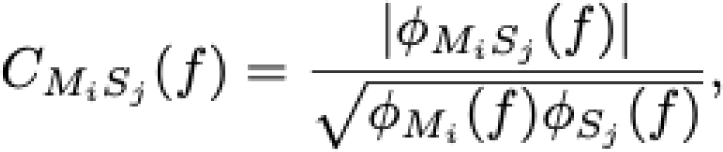

where ϕ (𝑓) is the cross-spectrum of the signals 𝑋 and 𝑌, while ϕ (𝑓) 𝑋𝑌 𝑋 and ϕ (𝑓) 𝑌 are their respective power spectral densities (PSD). We then average each one of this quantity across all modulator-sender pairs and across the two animals, in order to obtain the average coherence magnitude . The *sender-modulator-region* coherence is computed analogously, for each electrode in the modulator’s region (which is not a modulator itself) we compute the coherence with the sender. Similarly, the *sender-other-region* coherence is computed by calculating the mean coherence between electrodes which are not in modulators’ region and the sender site.

### Sender-Modulator coherence as a function of modulator ranking

We ranked modulators using the AUC from the ROC analysis of ‘hit’ and ‘miss’ events of the receiver’s response to the sender’s single-pulse stimulus (Fig. 2c). Sender-modulator-site coherence is then computed as a function of the modulator-ranking (from top ranked to bottom). For the computation of *sender-modulator-region* coherence as a function of the modulator ranking we include all the electrodes in the modulators’ region corresponding to the fraction of ranked modulators (Fig. 5d**-e**). For instance, when we compute the *sender-modulator-region* coherence for the top 10% ranked modulators, we include all the electrodes in these 10% top modulators’ brain regions, except the modulators themselves, and we compute their coherence with the sender-site. For the computation of the *sender-other-region* coherence as a function of the modulator ranking, we include all the electrodes in the other regions with respect to the fraction of top ranked modulators considered (by excluding regions which are modulator regions themselves) and we compute the coherence between these electrodes and the sender-site (Fig. 5d**-e**).

### Modulator-Receiver coherence

Modulator-receiver coherence is calculated analogously to the sender-receiver coherence discussed in the “**Sender-Modulator coherence**” Method Section, with the difference that the receiver site is used (instead of the sender-site) for the computation of the coherence with the modulator-site, the modulator-region, and all-other-regions (Fig. 6d**-e**)

### Receiver-Modulator coherence as a function of modulator ranking

Receiver-modulator coherence is calculated as a function of modulator ranking analogously to the Sender-Modulator coherence as a function of the modulator ranking discussed above.

### Modulator-Modulator theta-coherence network

To construct the modulator-modulator theta-coherence network shown in Fig. 8 we consider the sender-receiver pair with the largest number of modulators in each monkey, in order to investigate the largest possible modulator-modulator (MM) network available from our data (18 modulators for Monkey M, 14 modulators for Monkey A - see Fig. 8a and **Supp. Fig. S2a** respectively). However, the same analysis can be performed for any sender-receiver pair with any number of associated modulators. For each modulator pair we compute the modulator-modulator mean theta-coherence (averaged across the \theta-band frequency range). For a pair of modulators (M_i_, M_j_), this calculation results in the entry (i,j) in the matrix of Fig. 8b. In formulas, this quantity is computed as the mean of over the theta-frequency range, i.e. [4 - 10] Hz, with given by

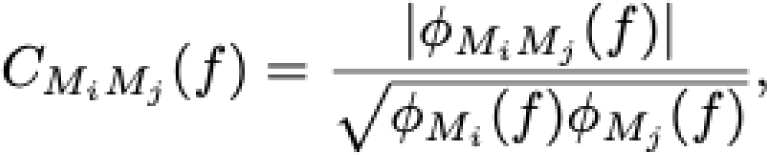

where ϕ (𝑓) is the the cross-spectrum of the signals of modulator M and M, while ϕ (𝑓) and ϕ (𝑓) are their respective power spectral densities. To quantify the significance of the modulator-modulator theta-coherence matrix entries we compute an analogous coherence for control electrodes, named modulator-control (MC) coherence, or 𝑀 𝐶 when referring to a pair of modulators M and M. In brief, 𝑀 𝐶 is the average between the M_i_C and M_j_C where M_i_C coherence between the modulator M_i_ and all the electrodes in the same region as the modulator M_j_ which are not modulators themselves for the same SR-pair. M_j_C coherence is computed analogously. In formulas M_i_C is:

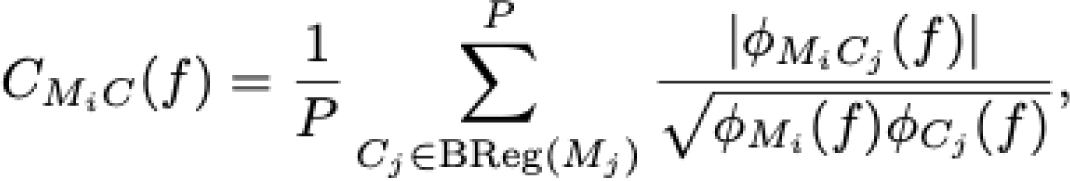

where the sum extends to all the P controls C_j_ in the same brain region as the modulator M_j_ (denoted as BReg(M_j_)), i.e. all the electrodes not modulating the same SR-pair in the same brain region. Finally 𝑀 𝐶 is the average between the M_i_C and M_j_C (Fig. 8c). We use the entries of 𝑀 𝐶 to construct a null distribution for the modulator-modulator coherence and compute the p-value theta-coherence matrix of Fig. 8d, where each (i,j) entry refers to the p-value associated with the M_i_ M_j_ coherence. Finally, we corrected this matrix for multiple comparison (FDR procedure ^52^ with α =0.05, Fig. 8e). The network diagram illustration of Fig. 8e is presented in Fig. 8f. In this diagram, a link between a pair of modulators exists if there exists the corresponding matrix entry in Fig. 8e, i.e. if the FDR corrected p-value for the modulator pair M_i_M_j_ is P_FDR_ < 0.05.

## Author contributions

G.D.F., S.Q., and B.P. conceived the study. G.D.F. and B.P. designed and implemented the coherence analysis. S.Q. and B.P. designed and implemented the experiments. S.Q., J.I.S., and B.P. performed the experiments and data collection. G.D.F and B.P. wrote the manuscript. G.D.F., B.P., and S.Q. revised the manuscript. B.P. supervised the work.

## Supplementary Information

**Table S1.** Type of brain connection for the brain regions relative to identified sender-receiver pairs identified via electrical stimulation. Brain connections are classified as direct/indirect or likely direct/indirect. The corresponding reference for each of these connections is provided.

| N. | Sender-Receiver | Type of connection | Reference |
| --- | --- | --- | --- |
| 1 | OFC - M1 | Likely indirect | 53 |
| 2 | OFC - ACC | Likely direct | 53 |
| 3 | CN - CN | Local |  |
| 4 | OFC - dPFC | Likely direct | 53 |
| 5 | CN - M1 | Likely indirect. Only direct if retrograde stimulation response | 10 |
| 6 | CN - ACC | Likely indirect. Only direct if retrograde stimulation response | 10 |
| 7 | CN - dPFC | Likely indirect. Only direct if retrograde stimulation response | 10 |
| 8 | CN - S1 | Likely indirect. Only direct if retrograde stimulation response | 10 |
| 9 | CN - OFC | Likely indirect. Only direct if retrograde stimulation response | 53 |
| 10 | CN - Put | Local | 53 |
| 11 | OFC - SMG | Likely direct | 53 |
| 12 | Amyg-OFC | Likely direct | 53 |
| 13 | Amyg-ACC | Likely direct | 54 |

**Table S2.** Number of modulators for each SR-pair, by Monkey, and type of SR-connection. Each row in the table reports one identified Sender-Receiver pair with the anatomical location of the Sender and Receiver site for such pair; whether the connection is direct (D), likely direct (LD), indirect (I), or likely indirect (LI); the number of identified modulators for the identified SR pair. Results are reported separately for monkey M and monkey A.

| **SR pair #** | **S region** | **R region** | **Type of connection** | **# of modulators** |
| --- | --- | --- | --- | --- |
| 1 | OFC | CN | D/I | 3 |
| 2 | OFC | CN | D/I | 3 |
| 3 | OFC | CN | D/I | 1 |
| 4 | OFC | CN | D/I | 3 |
| 5 | OFC | CN | D/I | 1 |
| 6 | OFC | CN | D/I | 11 |
| 7 | OFC | CN | D/I | 1 |
| 8 | OFC | dPFC | LD | 1 |
| 9 | OFC | M1 | LI | 3 |
| 10 | OFC | M1 | LI | 3 |
| 11 | OFC | ACC | LD | 1 |
| 12 | OFC | ACC | LD | 1 |
| 13 | OFC | ACC | LD | 2 |
| 14 | CN | M1 | D/I | 1 |
| 15 | CN | M1 | D/I | 13 |
| 16 | CN | M1 | D/I | 18 |
| 17 | CN | CN | LD | 12 |
| 18 | CN | CN | LD | 10 |
| 19 | CN | CN | LD | 4 |
| 20 | CN | CN | LD | 5 |
| 21 | CN | ACC | D/I | 1 |
| 22 | CN | ACC | D/I | 8 |
| 23 | CN | dPFC | D/I | 1 |
| 24 | CN | dPFC | D/I | 3 |

**Table S2. Number of modulators for each SR-pair, by Monkey, and type of SR-connection.**
| SR pair # | Sender region | Receiver region | Type of connection | # of modulators |
| --- | --- | --- | --- | --- |
| 1 | CN | ACC | D/I | 4 |
| 2 | CN | ACC | D/I | 14 |
| 3 | CN | S1 | D/I | 4 |
| 4 | CN | S1 | D/I | 7 |
| 5 | CN | OFC | D/I | 2 |
| 6 | CN | OFC | D/I | 3 |
| 7 | CN | CN | LD | 9 |
| 8 | CN | Put | LD | 1 |
| 9 | OFC | SMG | LD | 5 |
| 10 | OFC | ACC | LD | 2 |
| 11 | Amyg | OFC | LD | 3 |
| 12 | Amyg | ACC | LD | 7 |

**Table S3.** Modulator brain region and number of identified modulators in each region. Each row lists the number of modulators identified in each brain region for Monkey M and A separately.

| Region # | Modulator region | # of modulators |
| --- | --- | --- |
| 1 | CN | 27 |
| 2 | M1 | 52 |
| 3 | ACC | 4 |
| 4 | dPFC | 19 |
| 5 | OFC | 7 |

| Region # | Modulator region | # of modulators |
| --- | --- | --- |
| 1 | SMG | 4 |
| 2 | SPL | 16 |
| 3 | pCun | 4 |
| 4 | Put | 2 |
| 5 | ACC | 6 |
| 6 | M1 | 11 |
| 7 | CN | 4 |
| 8 | S1 | 5 |
| 9 | OFC | 5 |
| 10 | dPFC | 2 |
| 11 | BFB | 1 |
| 12 | Amyg | 1 |

**Figure S1.**
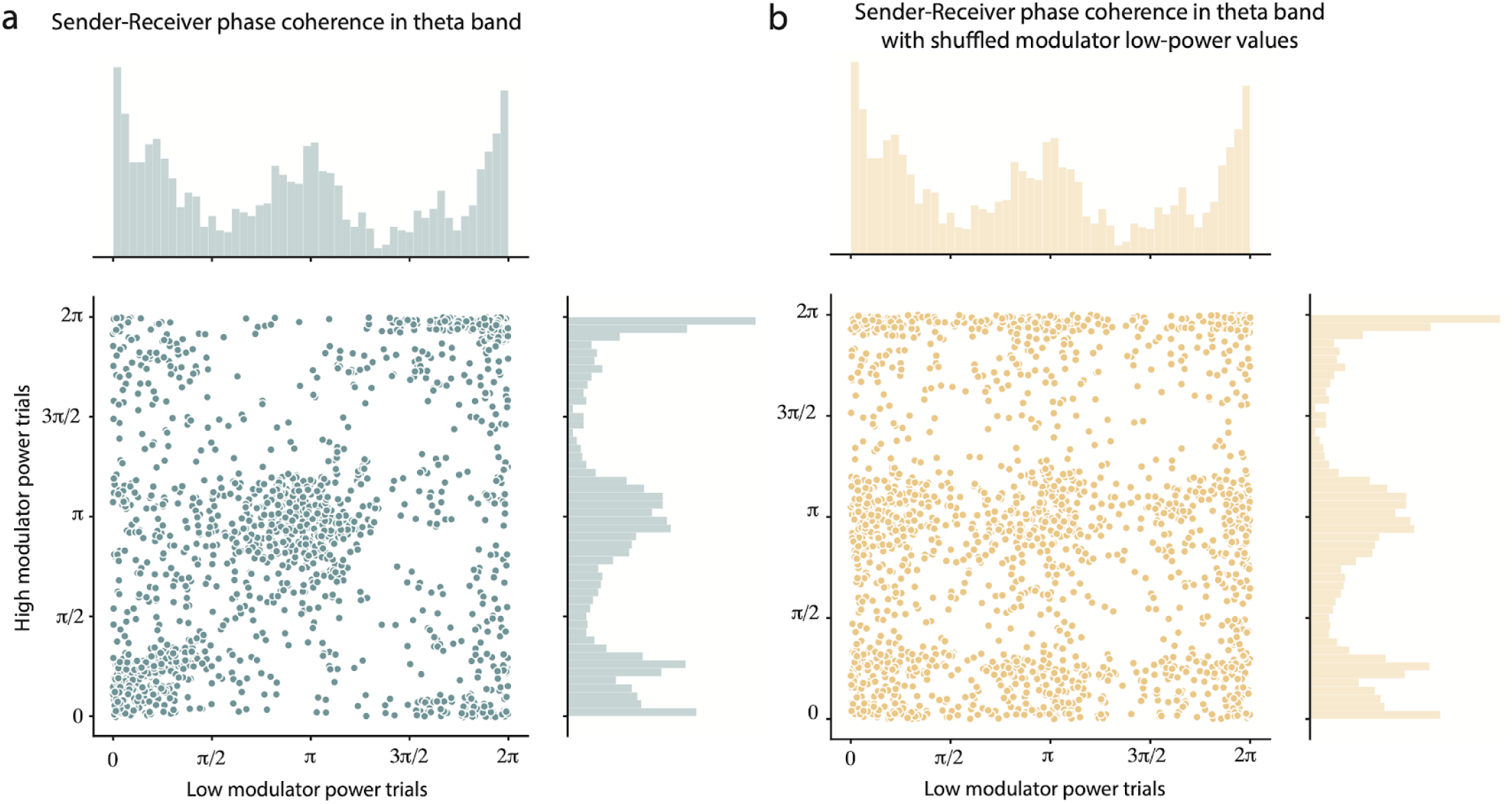
Theta-band Sender-Receiver coherence phase. **(a)** Scatter plot for the theta-band (4-10 Hz) coherence phase for epochs with high theta power in the identified modulator sites (vertical axis) and low theta power in the identified modulator sites (horizontal axis). The histogram along the horizontal axis presents the distribution of sender-receiver theta coherence phase during high modulator power epochs. The histogram along the vertical axis presents the distribution of sender-receiver theta coherence phase during low modulator power epochs. Sender site SR theta coherence is in-phase and out-of-phase when the modulator power is high and low. This result cannot be explained by volume conduction from the modulator sites. **(b)** Same as in **(a)** with shuffled low modulator power trials. The shuffling does not affect the histograms but it affects the scatter plots, showing that the distribution of values shown in the scatter plot in (a) is not a random effect.

**Figure S2.**
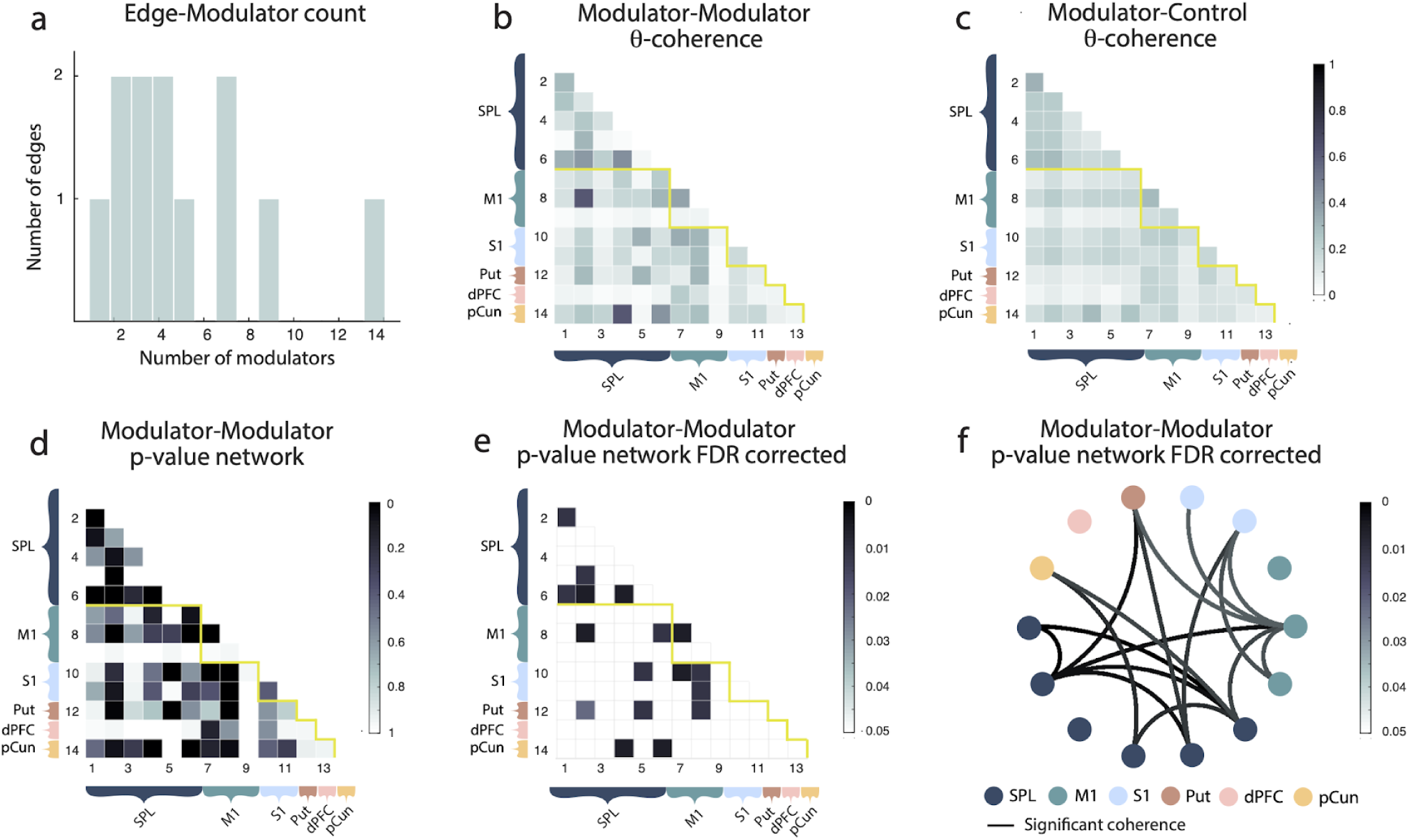
Network of modulators. **(a)** Empirical distribution of the number of SR-pairs with a given number of modulators. **(b)** Modulator-modulator pairwise theta-coherence for the session with the largest number of modulators (14 sites) for this monkey. Electrodes are organized by brain region, as illustrated in the panel. **(c)** Modulator-Control pairwise theta-coherence for the same session considered in (b) used to create a null-distribution for the modulator-modulator theta-coherence. **(d)** Statistical significance (p-value) of the modulator-modulator theta-coherence shown in panel **(a)**. **(e)** Same as in **(d)** after FDR correction for multiple comparison, values represent P_FDR_ . **(f)** Network representation of modulator-modulator statistical significant theta-coherence (network representation of the matrix shown in **(e)**: the existence of a link represents a statistical significant theta-coherence among a pair of modulators). All results refer to Monkey A.

